# Temporal orchestration of transcriptional and epigenomic programming underlying maternal embryonic diapause in a cricket model

**DOI:** 10.1101/2025.08.04.668555

**Authors:** Kosuke Kataoka, Yuta Shimizu, Ryuto Sanno, Yuichi Koshiishi, Ken Naito, Kei Yura, Toru Asahi, Shin G. Goto

**Author notes:** These authors equally contributed to this work (Kosuke Kataoka, Yuta Shimizu).

## Abstract

Maternal perception of environmental conditions can direct offspring developmental trajectories, providing adaptive flexibility across taxa. In the band-legged ground cricket *Dianemobius nigrofasciatus*, maternal exposure to short days induces embryonic diapause at the cellular blastoderm stage in offspring. Here, we investigate molecular mechanisms underlying this transgenerational adaptation through genome assembly (1.45 Gbp) and time-series transcriptomic analyses of diapause and non-diapause eggs from 12 to 72 hours post-oviposition. Despite morphological similarity, diapause-destined eggs show early upregulation of ATP-dependent chromatin remodeling genes at 24 hours. ATAC-seq reveals reduced chromatin accessibility at neural and cell cycle-related genes. Time-series clustering identifies precocious shifts in RNA processing machinery (peaking at 24 versus 40 hours in non-diapause eggs), followed by metabolic regulation toward amino acid catabolism and gluconeogenesis sustaining long-term survival during developmental arrest. Our findings reveal diapause as actively coordinated molecular programming involving epigenetic, transcriptional, and metabolic remodeling, providing insights into transgenerational environmental adaptation.

## Introduction

Across the tree of life, organisms have evolved remarkable strategies to survive temporarily hostile environments through dormancy, a physiological state characterized by dramatically reduced metabolic activity that enhances survival during adverse conditions ^1,2^. Among the most widespread environmental challenges is seasonality, which poses significant physiological demands on organisms inhabiting temperate regions. To survive periods unsuitable for growth and reproduction, diverse taxa have independently evolved dormancy strategies: hibernation and torpor in mammals and birds, as well as embryonic developmental arrest in vertebrates and developmental pauses in numerous invertebrates, notably insects ^3^.

Among these strategies, diapause emerges as a programmed form of dormancy employed by many insect species in anticipation of seasonal stressors. Diapause can occur at any developmental stage—egg, larva, pupa, or adult—depending on the species-specific life cycle strategy ^4^. A number of common physiological events are associated with diapause, including metabolic suppression and developmental arrest ^5^, enhanced resistance to cold and desiccation ^6,7^, and accumulation of energy reserves ^8^.

In many species, individuals enter diapause upon receiving a token stimulus. However, a particularly intriguing mechanism involves maternal influence, whereby the mother receives environmental signals and transfers this information to her progeny. This transgenerational effect is referred to as the maternal effect ^9^. Maternal diapause induction is particularly valuable for embryonic diapause, a situation in which an individual has limited access to or time to independently evaluate seasonal conditions. This transgenerational adaptation has been well-documented across diverse taxa, including mites ^10^, crickets ^11–13^, rice leaf bug ^14^, flies ^15^, mosquitoes ^16^, and parasitic wasps ^17,18^.

Despite the growing interest in maternal diapause induction, the underlying molecular mechanisms remain poorly understood. Recent omics studies have highlighted the importance of alterations in endocrine signaling, cell proliferation, metabolism, and energy production in maternal diapause induction ^19,20^; however, these investigations have not comprehensively addressed how maternal influences impact specific molecular pathways initiating diapause, due to limited sampling periods. A comprehensive understanding of the molecular processes through which maternal environmental signals redirect the developmental trajectory towards diapause requires investigation of multiple time points throughout early development.

*Dianemobius nigrofasciatus*, the band-legged ground cricket, serves as a prominent model for investigating the maternal effect on embryonic diapause (Fig. 1a, b) ^13,21^. Adult females reared under long-day conditions produce eggs that develop directly into nymphs without undergoing diapause (hereafter called long-day eggs), whereas those exposed to short-day conditions lay diapause-destined eggs (hereafter called short-day eggs) (Fig. 1b) ^12,13^. A previous morphological study ^22^ established that long- and short-day eggs remain visually indistinguishable until 56 hours post-oviposition (hpo). During the first 40 hpo, energids (early embryonic cells) proliferate and migrate to the egg surface in both egg types. Subsequently, between 40 and 56 hpo, both types of eggs undergo cellular blastoderm formation. However, by 72 hpo, a clear morphological divergence emerges: long-day eggs proceed to form a germ band at the posterior pole, while short-day eggs arrest at the cellular blastoderm stage without germ band formation. Although the involvement of suppressed cell cycle regulators, small silencing RNAs, and segment patterning genes has been suggested ^23^, the precise molecular mechanisms underlying maternal diapause induction remain unknown.

**Figure 1.**
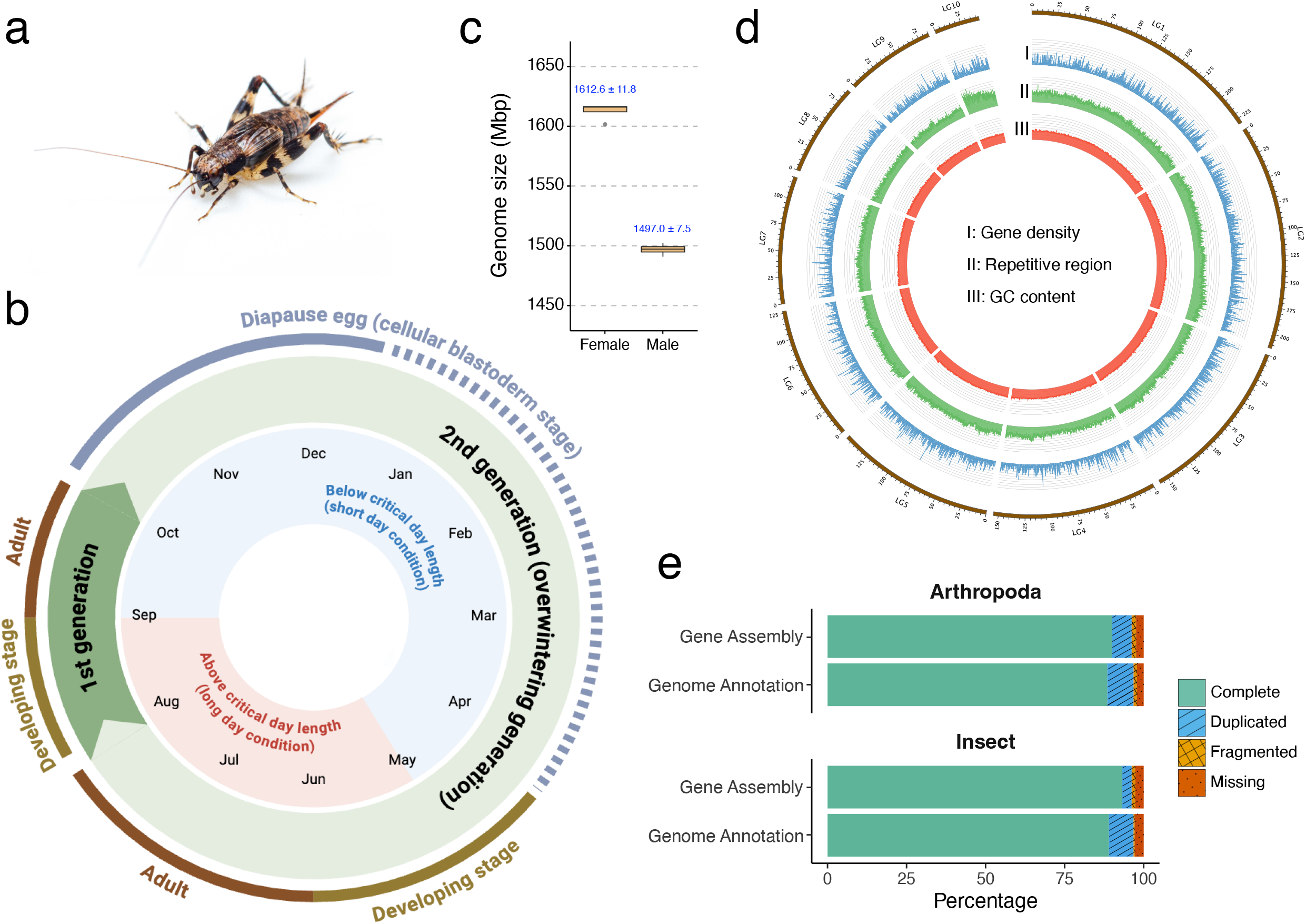
Life cycle and genomic characterization of the band-legged ground cricket, *Dianemobius nigrofasciatus*. **(a)** Adult female *D. nigrofasciatus*. **(b)** Annual life cycle diagram in Osaka, Japan, illustrating photoperiod-dependent developmental transitions and diapause regulation. The outer ring represents the developmental stages (egg, nymph, and adult) and the diapause egg state. The inner circle shows seasonal progression with photoperiod-dependent phases: the pink area indicates the period with the longer critical day length, whereas the blue area indicates the period with the shorter critical day length. The first-generation adults emerge in September-October under short-day conditions and produce predominantly diapause-destined eggs that overwinter in the cellular blastoderm stage, whereas the second-generation adults appear in June-August under long-day conditions and produce non-diapause eggs. **(c)** Flow cytometric analysis of brain cells from adult females and males showing genome size estimation. Box plots show the distribution of genome size measurements, with blue numbers indicating mean ± 95% confidence interval for each sex. **(d)** Circos plot depicting genomic features across the ten linkage groups (LG1-10). From outer to inner tracks: gene density (I), repetitive element content (II), and GC content distribution (III). **(e)** BUSCO completeness assessment of the genome assembly and gene annotation against Arthropoda and Insecta databases via the gVolante web server. Colors indicate complete single-copy (solid green), completely duplicated (blue with diagonal stripes), fragmented (yellow crosshatched pattern), and missing (red with black dot pattern) genes.

In this study, we assembled a high-quality genome of *D. nigrofasciatus* and conducted a comprehensive time-series transcriptomic analyses of long- and short-day eggs to elucidate the molecular events occurring before and after the onset of embryonic diapause. Based on the morphological milestones, we strategically selected timepoints that would capture molecular changes before (12 and 24 hpo), during (40 and 56 hpo), and after (72 hpo) the critical developmental decision point where diapause and non-diapause trajectories diverge. Our findings suggest that this process is orchestrated not by a straightforward developmental switch, but rather by a sophisticated temporal orchestration of molecular programs.

## Results

### Chromosome-scale genome assembly and gene annotation of *D. nigrofasciatus*

To investigate the molecular mechanisms of maternal diapause induction, we generated a chromosome-scale genome assembly of *D. nigrofasciatus* using short-read, long-read, and Omni-C sequencing technologies. Flow cytometry revealed genome size differences between sexes, with males (1496.9 Mbp) having smaller genomes than females (1612.6 Mbp) (Fig. 1c; Supplementary Table 1), consistent with the XO/XX sex determination system in crickets ^24^. *K*-mer analysis estimated a haploid genome size of 1.24 Gbp, which is smaller than the flow cytometry estimates, likely due to underrepresentation of repetitive sequences in short-read data (Supplementary Fig. 1; Supplementary Table 2).

The hybrid assembly strategy integrated 100.84× Illumina, 19.87× Oxford Nanopore, and 78.82× Omni-C coverage (Supplementary Table 3), resulting in a 1.45 Gbp assembly across 482 scaffolds with a scaffold N50 of 154 Mbp and minimal contamination (Supplementary Fig. 2). 87.1% of sequences were anchored into 10 linkage groups (Fig. 1d; Supplementary Fig. 3; Supplementary Tables 4, 5). Coverage analysis of male genomic sequences revealed that LG8 and LG9 constitute the X chromosome (Supplementary Fig. 4; Supplementary Table 6).

Assembly quality assessment using BUSCO (Benchmarking Universal Single-Copy Orthologs) ^25^ demonstrated high completeness, with 90.0% complete single-copy and 6.2% duplicated orthologs for the Arthropoda dataset (n=1,013) (Fig. 1e; Supplementary Table 4). When evaluated against the more specific Insecta dataset (n=1,367), the assembly showed even higher completeness with 93.2% single-copy and 3.0% duplicated orthologs (Fig. 1e; Supplementary Table 4). Gene annotation using BRAKER3 ^26–28^, StringTie2 ^29^, and GeMoMa ^30^ yielded 20,812 protein-coding genes (Supplementary Table 7). Phylogenetic analysis of 334 single-copy orthologs placed *D. nigrofasciatus* within Trigonidiidae, sister to Gryllidae (Supplementary Fig. 5),consistent with previous studies ^31–33^.

This high-quality genome assembly and gene annotation provide crucial resources for elucidating the mechanisms of transgenerational diapause programming.

### Gradual molecular divergence in maternal diapause induction

To elucidate the molecular mechanisms underlying maternal diapause induction, we performed comprehensive time-series RNA-seq analysis of both long- and short-day eggs at five developmental timepoints: 12, 24, 40, 56, and 72 hpo (Fig. 2a). Principal component analysis (PCA) of the transcriptome data revealed intriguing patterns in the temporal dynamics of developmental divergence (Fig. 2b). Given that maternal diapause induction presumably involves signaling from mother to offspring, we initially expected substantial transcriptomic differences even at the earliest timepoint. However, molecular differences between long- and short-day eggs were minimal at 12-24 hpo, despite both egg types having been exposed to different maternal environments. The transcriptomic distance between the two egg types then expanded progressively from 24 to 72 hpo, with the most dramatic separation occurring between 56 and 72 hpo, coinciding with morphological divergence.

Differential expression analysis substantiated these observations, revealing a progressive increase in the number of differentially expressed genes (DEGs) between long- and short-day eggs over time (|log_2_ fold change| > 1, adjusted *p*-value < 0.05) (Fig. 2c, left panel; Supplementary Table 8). Remarkably, at 12 hpo, merely 31 genes showed differential expression (16 upregulated, 15 downregulated in short-day eggs). This number increased modestly to 120 DEGs at 24 hpo, then expanded more substantially to 371 DEGs at 40 hpo, 486 DEGs at 56 hpo, and culminated in 1,994 DEGs at 72 hpo (1,078 upregulated, 916 downregulated in short-day eggs).

**Figure 2.**
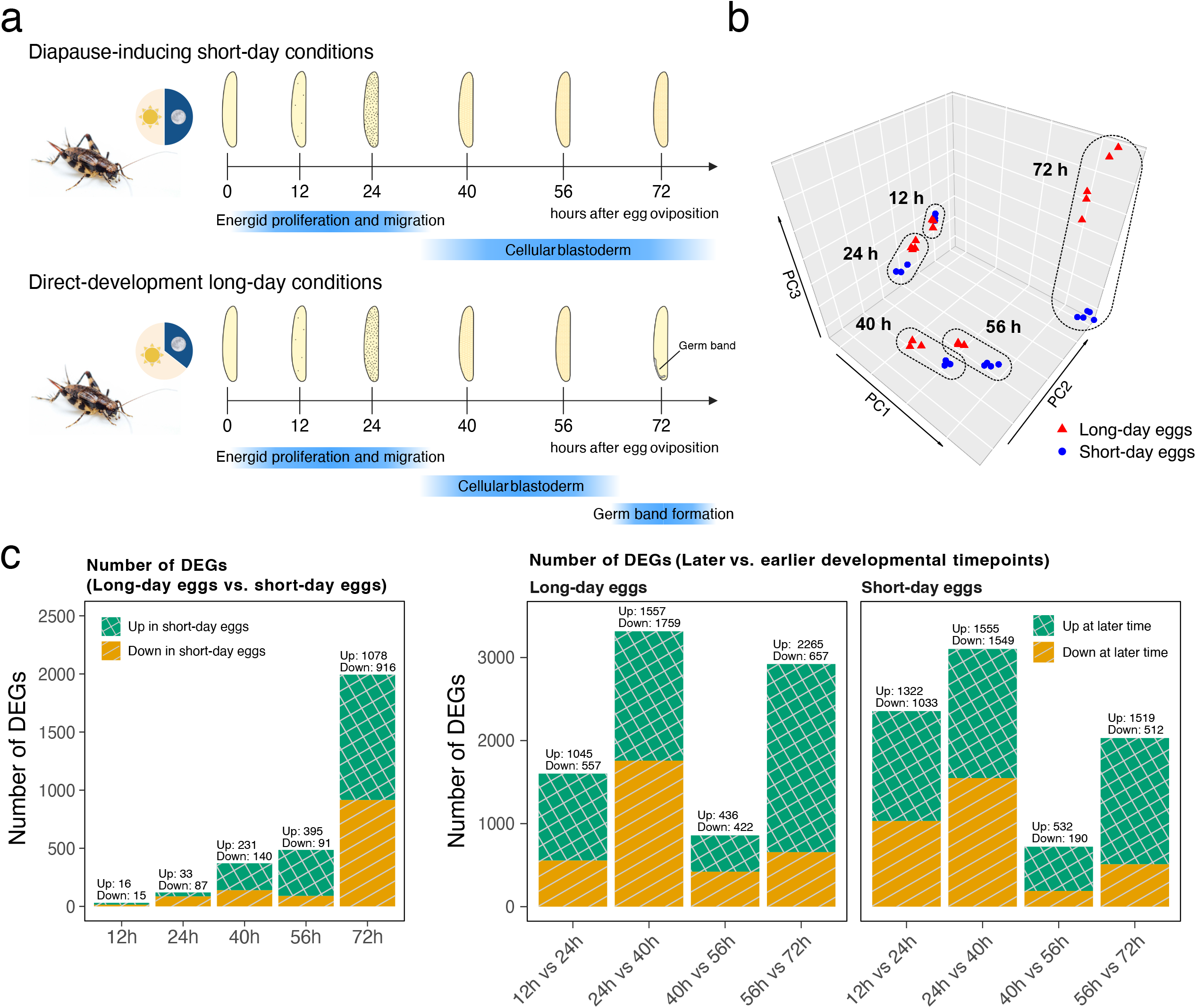
Transcriptome dynamics during early embryogenesis in long- and short-day eggs of *D. nigrofasciatus*. **(a)** Schematic representation of developmental progression from 0 to 72 hours post-oviposition (hpo) at 25°C. Adults reared under short-day conditions produce diapause-destined eggs (short-day eggs), while those under long-day conditions produce eggs that undergo direct development (long-day eggs). The morphological features of early embryonic stages were characterized by Tanigawa et al. (2009). In both egg types, nuclei migrate from the yolk mass to the egg surface by 24 hpo. Cellular blastoderm formation occurs between 40-56 hpo in both trajectories. While long-day eggs proceed to form a germ band by 72 hpo, short-day eggs arrest development at the cellular blastoderm stage, maintaining this state throughout the diapause period. **(b)** Three-dimensional principal component analysis of transcriptome profiles across developmental time points. Red and blue marks indicate long- and short-day eggs, respectively. Dashed line circles indicate clustering of samples corresponding to each developmental time point. **(c)** *Left panel*: Number of differentially expressed genes (DEGs) between long- and short-day eggs at each developmental timepoint (12, 24, 40, 56, and 72 hpo). DEGs were defined as genes with |log_2_ fold change (FC)| > 1 and adjusted *p*-value < 0.05. Green bars with crosshatched pattern indicate genes upregulated in short-day eggs, while yellow bars with diagonal stripes represent genes downregulated in short-day eggs. *Right panels*: Temporal dynamics of gene expression changes within each egg type, comparing later versus earlier developmental timepoints. For long- (left) and short-day eggs (right), bars show the number of genes upregulated (green with crosshatched pattern) or downregulated (yellow with diagonal stripes) at the later timepoint relative to the earlier timepoint in each comparison (|log_2_ FC| > 1, adjusted *p*-value < 0.05).

Analysis of temporal gene expression changes between consecutive timepoints revealed three distinct developmental phases (Fig. 2c, right panel; Supplementary Table 9). The first major transcriptional transition (24-40 hpo) coincided with energid proliferation and migration, showing 3,316 DEGs in long-day eggs and 3,104 DEGs in short-day eggs. Transcriptional activity then decreased during cellular blastoderm formation (40-56 hpo), with 858 and 722 DEGs in long- and short-day eggs, respectively. The final phase (56-72 hpo) showed divergent patterns, particularly in long-day eggs (2,922 DEGs) compared to short-day eggs (2,031 DEGs), reflecting the initiation of germ band formation in long-day eggs while short-day eggs undergo molecular changes associated with diapause ^22^.

### Temporal coordination and divergence of gene expression patterns reveal diapause preparation dynamics

To comprehensively characterize these temporal dynamics and identify genes with similar expression patterns across developmental time, we performed fuzzy c-means clustering using Mfuzz ^34^ on the time-series transcriptome data. This soft clustering assigns genes to multiple clusters with varying membership degrees. This feature makes it particularly suitable for capturing the continuous nature of developmental processes. Following independent clustering of each dataset (long- and short-day eggs), we identified cluster pairs between the two egg types that showed significant gene overlap using upper-tail hypergeometric tests (Fig. 3a; Supplementary Fig. 6, 7). These overlapping cluster pairs were further classified based on their temporal expression patterns: “homochronic clusters” displayed synchronized expression dynamics between egg types, while “heterochronic clusters” exhibited temporal shifts in peak expression timing, revealing potential differences in developmental timing between the two trajectories (Supplementary Fig. 8; Supplementary Table 10).

To understand which developmental processes remain conserved, we first examined homochronic gene clusters that maintained synchronized expression between egg types. The majority of homochronic cluster pairs exhibited peak expression during the early developmental periods (12-24 hpo), suggesting that fundamental embryonic processes are largely conserved before the diapause trajectory diverges (Supplementary Fig. 9; Supplementary Table 11).

**Figure 3.**
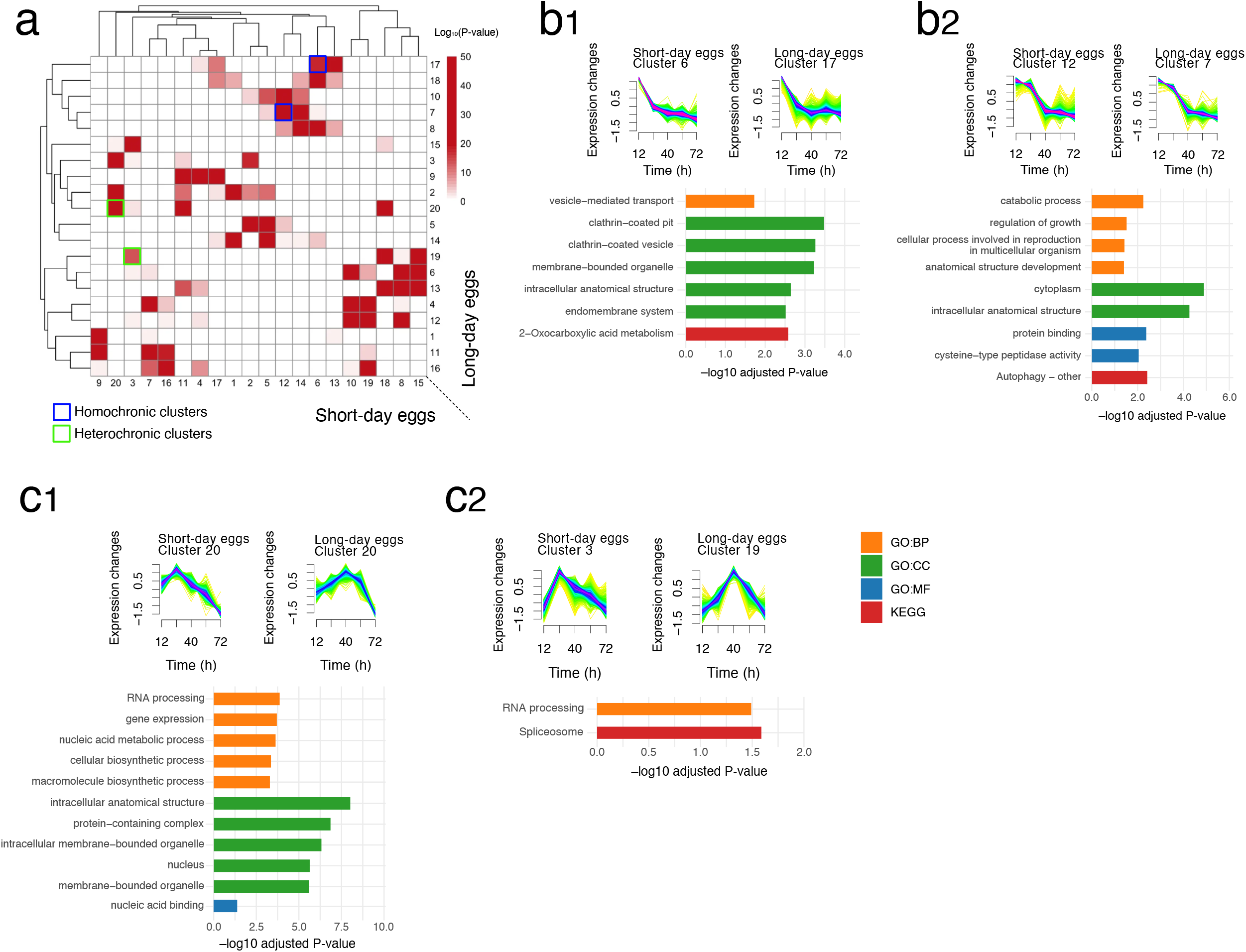
Comparative analysis of temporal gene expression dynamics between long- and short-day eggs. **(a)** Heat map showing the significance of gene overlap between temporal expression clusters in long- and short-day eggs across five developmental timepoints (12, 24, 40, 56, and 72 hpo) as derived from an upper-tail hypergeometric test. Gene expression profiles were clustered using the Mfuzz soft clustering algorithm. Numbers on the right and bottom axes represent cluster IDs for long-day and short-day eggs, respectively. Color intensity represents −log_10_(adjusted *p*-value), with darker red indicating more significant overlap between clusters. Homochronic clusters (highlighted in blue boxes) show conserved temporal expression patterns between the two egg types. Heterochronic clusters (highlighted in green boxes) exhibit shifted temporal dynamics between long- and short-day eggs. **(b1, b2, c1, and c2)** Expression trajectories show representative homochronic **(b)** or heterochronic **(c)** cluster pairs, where color intensity reflects membership scores calculated by Mfuzz (magenta indicating high membership and green indicating low membership). Bar graphs display functional enrichment analysis results using overlapping genes, with terms categorized as GO Biological Process (GO:BP, orange), GO Cellular Component (GO:CC, green), GO Molecular Function (GO:MF, blue), or KEGG pathways (red). For each category, the top five most significantly enriched terms (ranked by adjusted *p*-value) are shown; when fewer than five terms were significant, all enriched terms are displayed. Bar length represents −log_10_(adjusted *p*-value).

Analysis of these early-peaking homochronic clusters revealed common activation of essential cellular machinery: genes encoding components of clathrin-coated vesicles and membrane trafficking systems showed high initial expression followed by continuous decline, likely reflecting the early embryo’s use of maternally supplied membrane and vesicle pools required for cellularization, together with maternally provided trafficking proteins and maternally deposited mRNAs that are translated during early development (Fig. 3b) ^35–37^. Both egg types also showed coordinated activation of autophagy-related genes, peptidases, and catabolic processes at 12-24 hpo, suggesting a conserved program for recycling maternal components during early embryogenesis (Fig. 3b) ^37–40^.

In contrast, heterochronic clusters unveiled critical timing differences in key molecular pathways (Supplementary Fig. 10; Supplementary Table 12). Notably, genes involved in RNA processing and spliceosome function exhibited marked temporal advancement in short-day eggs (Fig. 3c). While these genes peaked at 40 hpo in long-day eggs, corresponding to the onset of cellular blastoderm formation, they reached maximum expression already at 24 hpo in short-day eggs. This precocious activation of RNA processing machinery coincides with the upregulation of chromatin remodeling factors, as shown in the subsequent epigenetic analyses (Fig. 4a, b), suggesting a coordinated early establishment of the molecular foundation required for developmental arrest.

**Figure 4.**
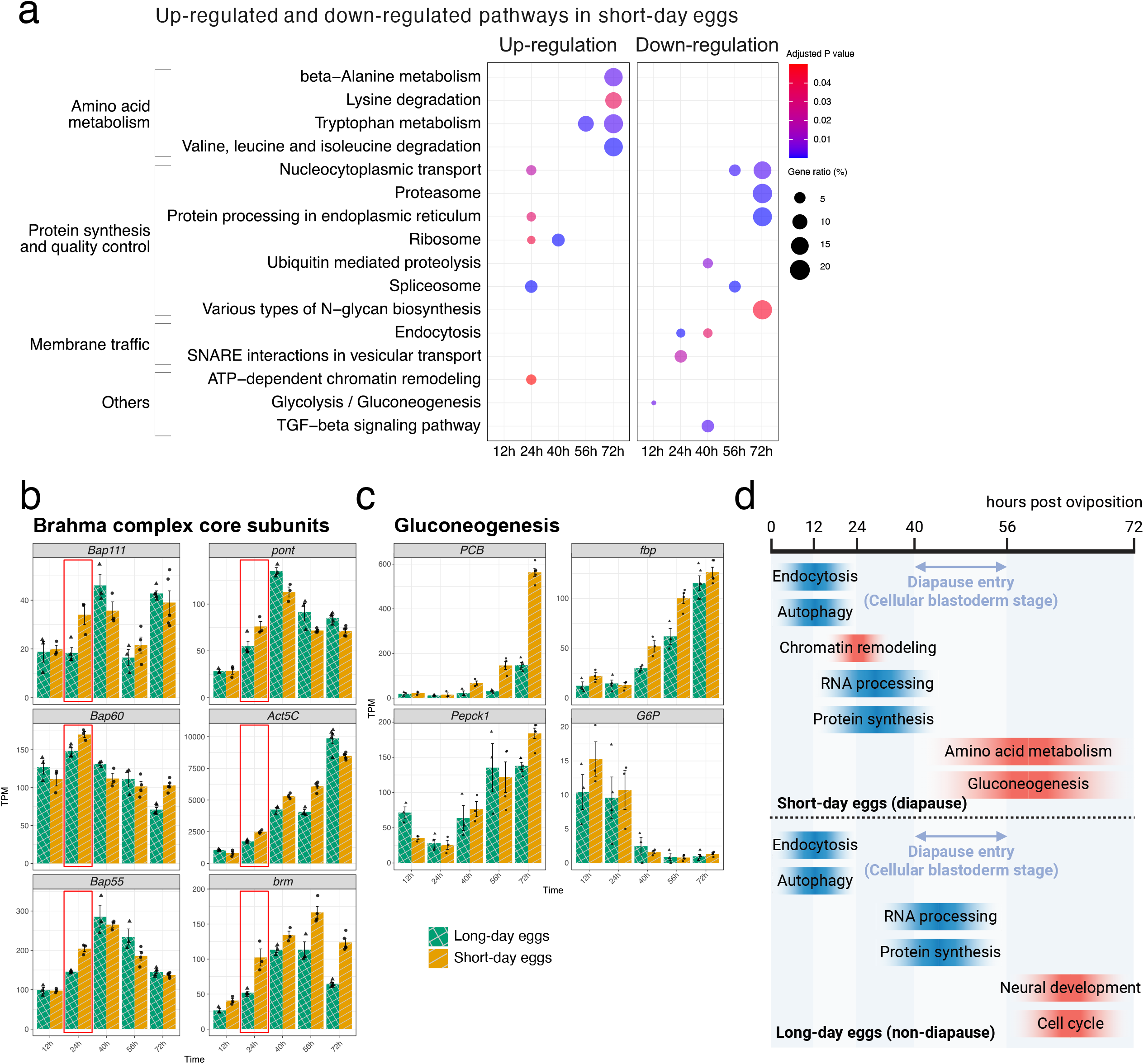
Temporal coordination of biological processes during early embryogenesis in long- and short-day eggs. (**a)** KEGG pathway enrichment analysis of differentially expressed genes (DEGs) identified between long-day and short-day conditions at each time point (adjusted *p*-value < 0.05). Dot size represents the percentage of DEGs annotated to each pathway, and color intensity indicates the adjusted *p*-value. **(b and c)** Temporal expression patterns (TPM values) of genes involved in chromatin remodeling **(b)** and gluconeogenesis **(c).** The red frames in (b) highlight the critical periods of expression divergence between long- and short-day eggs. **(d)** Schematic representation of key biological processes and their temporal dynamics from 0 to 72 hpo. Both long- and short-day eggs show similar timing of membrane trafficking processes (endocytosis and autophagy) during early embryogenesis (12-24 hpo). Chromatin remodeling occurs within a specific time window around 24 hpo. Prior to diapause entry at the cellular blastoderm stage, short-day eggs show advanced activation of RNA processing and protein synthesis compared to long-day eggs. During later development (>56 hpo), short-day eggs activate amino acid metabolism and gluconeogenesis pathways, while long-day eggs proceed with neural development and cell cycle progression. Blue-shaded processes indicate molecular programs that are activated in both short-day (diapause) and long-day (non-diapause) eggs, whereas red-shaded processes indicate pathways that are specifically activated in only one developmental trajectory.

### Early epigenetic remodeling drives diapause preparation

Differentially expressed genes unveiled the molecular mechanisms initiating diapause (Supplementary Table 13). At 12 hpo, the sole significantly enriched pathway was glycolysis/gluconeogenesis, showing downregulation in short-day eggs (Fig. 4a). However, by 24 hpo, this initial metabolic shift had expanded into comprehensive molecular programming, with multiple pathways emerging as significantly enriched between egg types. Most strikingly, ATP-dependent chromatin remodeling genes showed specific upregulation in short-day eggs at this timepoint (Fig. 4a). Detailed examination revealed increased expression of core subunits of the Brahma (SWI/SNF) complex, including *Bap111, pont*, and *Bap60* (Fig. 4b). The Brahma complex is a critical and conserved regulator of chromatin accessibility and gene expression ^41–43^, suggesting that epigenetic remodeling may serve as a molecular switch for diapause induction.

To test this hypothesis, we performed ATAC-seq analysis on 24 hpo eggs to assess genome-wide chromatin accessibility. Short-day eggs exhibited reduced chromatin accessibility specifically near genes associated with neural development and cell cycle regulation (Supplementary Fig. 11; Supplementary Table 14, 15). Notable examples include *Kr-h1*, encoding a juvenile hormone-responsive transcription factor essential for developmental progression ^44^, and *crol*, an ecdysone-inducible transcription factor that recruits Polycomb group proteins ^45,46^.

Additionally, chromatin closure was observed at *Tlk* (Tousled-like kinase), which triggers S-phase entry and histone assembly ^47^, and *Ssrp*, a FACT complex subunit required for transcription initiation ^48^. Neural signaling genes also showed reduced accessibility, including synaptic components such as *Tollo* (Toll-6) and *nAChRα1*. This selective chromatin closure likely contributes to the developmental arrest characteristic of diapause. The timing of these chromatin changes before morphological divergence suggests that epigenetic remodeling is not a consequence but rather a driver of diapause preparation.

### Metabolic regulation and developmental signaling orchestrate diapause preparation

Following the early establishment of epigenetic and transcriptional regulatory programs, a comprehensive metabolic and developmental restructuring emerged as the next major phase of diapause preparation. While initial changes were subtle, a dramatic transformation unfolded from 40 hpo onwards, intensifying through the morphological divergence point.

The metabolic shift was characterized by coordinated activation of multiple amino acid degradation pathways. By 56-72 hpo, valine, leucine, and isoleucine degradation, tryptophan metabolism, lysine degradation, and beta-alanine metabolism were significantly upregulated in short-day eggs (Fig. 4a). This was accompanied by increased expression of key gluconeogenic enzymes, with phosphoenolpyruvate carboxykinase 1 (*Pepck1*) showing marked upregulation from 40 hpo and pyruvate carboxylase (*PCB*) increasing progressively from 56 hpo onwards (Fig. 4c). Genes encoding malate-aspartate shuttle components (*Got2* and *Mdh1/2*) also showed elevated expression in short-day eggs at 56-72 hpo (Supplementary Fig. 12), suggesting enhanced mitochondrial-cytoplasmic metabolite exchange. These enzymes enable the conversion of amino acid-derived carbon skeletons into glucose ^8,49^, suggesting a metabolic strategy to maintain energy homeostasis during impending developmental arrest.

Complementing this metabolic regulation, protein homeostasis pathways showed coordinated adjustment. From 56 hpo onwards, coinciding with peak metabolic restructuring, proteasome, protein processing in the endoplasmic reticulum, and spliceosome pathways were significantly suppressed in short-day eggs (Fig. 4a). This comprehensive downregulation of both protein synthesis and degradation machinery effectively pauses the cellular protein economy to preserve resources for extended diapause.

Concurrent with these metabolic changes, key developmental signaling pathways diverged between diapause and non-diapause trajectories. At 40 hpo, we observed significant downregulation of TGF-beta signaling in short-day eggs (Fig. 4a). As a master regulator of cell proliferation and differentiation ^50,51^, suppression of TGF-beta signaling effectively applies molecular brakes on developmental progression, occurring before morphological divergence.

## Discussion

Our comprehensive molecular analysis reveals that maternal diapause induction in *D. nigrofasciatus* proceeds through a precisely orchestrated temporal cascade of molecular events (Fig. 4d). Both long- and short-day eggs initially share common early developmental processes, including endocytosis and autophagy for recycling maternal components (12-24 hpo). However, short-day eggs diverge through a distinctive molecular program: early activation of chromatin remodeling machinery and precocious RNA processing (24-40 hpo), followed by progressive metabolic restructuring with upregulation of amino acid catabolism and gluconeogenesis pathways that continues through 72 hpo. In contrast, long-day eggs maintain sustained protein synthesis machinery, RNA processing at normal developmental timing, and active cell cycle progression leading to cellular differentiation. This temporal progression demonstrates that diapause preparation is not a simple developmental arrest but rather an active programming process requiring substantial molecular investment before the morphological divergence point.

An important consideration in interpreting this temporal cascade is the timing of the maternal-to-zygotic transition (MZT), during which developmental control shifts from maternally deposited transcripts to embryo genome–derived transcription ^52^. Although the precise timing of MZT has not been formally characterized in *D. nigrofasciatus*, previous analyses in this species demonstrated clear temporal increases in multiple developmental regulators, including segmentation-related genes, between 12 and 40 hpo, suggesting the onset of zygotic transcription during this period ^23^. In line with these observations, our RNA-seq data show minimal photoperiod-dependent differences at 12 hpo, followed by progressive divergence from 24 hpo onward, suggesting that embryo genome transcription is already active by 12–24 hpo.

Under this framework, maternal diapause induction is unlikely to operate solely through differential deposition of maternal mRNAs. Instead, maternal photoperiodic experience likely establishes distinct epigenetic states in the egg, such as chromatin accessibility, that subsequently bias embryo-driven transcriptional programs toward diapause or direct development. A similar principle has been demonstrated in *Drosophila melanogaster*, in which MZT occurs well before cellularization and early chromatin and transcriptional states critically shape later developmental trajectories ^52^.

The temporal advancement of components of RNA processing in short-day eggs, peaking at 24 hpo versus 40 hpo in long-day eggs, represents a striking example of heterochronic gene expression. This precocious activation likely serves multiple adaptive functions. First, the early investment in RNA processing capacity could enable the preparation of stable mRNA pools that persist throughout the diapause period ^53,54^. Second, this enhanced RNA processing machinery may facilitate rapid transcriptional responses upon receiving diapause-terminating signals, allowing quick resumption of development ^53,54^. Third, it may establish diapause-specific alternative splicing programs that generate protein isoforms optimized for dormancy ^55^. The sustained elevation of these components throughout diapause preparation suggests they play a central role in maintaining the molecular readiness for eventual developmental resumption.

The early upregulation of Brahma complex components at 24 hpo, preceding any morphological changes, suggests that chromatin remodeling serves as the primary molecular switch for diapause induction. This finding aligns with emerging evidence that epigenetic mechanisms underlie phenotypic plasticity across diverse organisms ^56–59^. The selective reduction in chromatin accessibility at developmental regulators such as *Kr-h1* and *crol* effectively locks cells into a pre-differentiation state, preventing the normal progression of embryonic development ^44,45^. Both genes are central components of juvenile hormone (JH) and ecdysteroid signaling cascades, which are essential endocrine pathways governing insect growth and developmental progression.

Notably, however, our RNA-seq data did not reveal clear differences in juvenile hormone or ecdysteroid signaling pathways between long- and short-day eggs at 24 hpo. Consistent with this, prior physiological assays in *D. nigrofasciatus* did not detect egg JH III or photoperiod-dependent differences in egg 20-hydroxyecdysone during early development ^60^. These observations suggest that early chromatin closure at endocrine-responsive developmental regulators does not reflect active hormonal signaling at this stage, but rather a preemptive epigenetic mechanism that suppresses the embryo’s capacity to respond to later endocrine cues required for developmental progression. Within this framework, rather than being directly specified by maternally deposited Brahma transcripts, we propose that maternal photoperiodic perception establishes distinct epigenetic contexts in the egg, within which the Brahma complex might function as an executor that stabilizes and amplifies these maternally primed regulatory states. This epigenetic regulation occurs during a critical window (within 24 hpo), demonstrating how maternal environmental signals can be translated into stable epigenetic modifications that fundamentally alter offspring developmental trajectory. Intriguingly, this 24-hour window coincides with the period during which temperature pulses can still reverse diapause fate decisions ^61^, suggesting that chromatin remodeling may define developmental fate plasticity.

In *D. nigrofasciatus*, parental RNAi is technically feasible and could, in principle, enable functional validation of the Brahma complex ^62^. However, knockdown of Brahma has been reported to severely impair fertility and fecundity in other insect species, including coleopteran and hemipteran models ^63^, rendering such approaches impractical for maintaining sufficient egg production in this system. Importantly, diapause fate in *D. nigrofasciatus* eggs can be experimentally induced or prevented by temperature manipulation ^63^, and diapause termination occurs spontaneously after prolonged arrest and is accelerated by winter-like chilling ^64^.

Therefore, examining Brahma complex expression and chromatin states under experimentally altered diapause induction and termination trajectories provides a realistic and informative framework for assessing its functional relevance *in vivo*.

The progressive metabolic restructuring observed from 56 hpo onwards in short-day eggs reveals a sophisticated survival strategy distinct from simple developmental arrest. The coordinated upregulation of amino acid catabolism pathways and gluconeogenic enzymes indicates a fundamental shift from anabolic growth metabolism to a catabolic maintenance mode. This metabolic flexibility enables short-day eggs to utilize maternally deposited amino acids as an energy source while preserving essential proteins needed for eventual development ^53,54^. The timing of these changes, beginning before morphological divergence and intensifying through 72 hpo, indicates that metabolic programming is an integral preparatory component rather than a consequence of arrest. This preemptive metabolic adjustment, coupled with the suppression of protein synthesis and proteasome pathways, creates a molecular economy optimized for long-term survival while maintaining developmental competence ^13,21^.

The remarkable plasticity of *D. nigrofasciatus*, which can facultatively switch between diapause and direct development based on maternal photoperiod experience, makes it a unique model for understanding how environmental information is translated into developmental outcomes ^64,65^.

Our findings illuminate how subtle maternal signals can be progressively amplified into dramatically different developmental outcomes through self-reinforcing molecular cascades that combine epigenetic, transcriptional, and metabolic regulation. This temporal orchestration framework likely extends beyond insects to other organisms that employ maternal programming of developmental dormancy, including delayed implantation in mammals ^65,66^, suggesting that coordinated molecular timing represents a fundamental mechanism for transgenerational environmental adaptation across diverse organisms. This cricket model system provides a powerful framework for understanding the universal principles governing how organisms across the tree of life integrate environmental information to orchestrate developmental decisions through conserved molecular mechanisms.

## Methods

### Sample collection

The *D. nigrofasciatus* individuals used in this study was collected in a grassy field on the campus of Osaka Metropolitan University (34°35′N, 135°30′E) and the bank of the Yamato-gawa River (34°35′N, 135°30′E), Japan, from June to November 2022 and 2023. The rearing temperature (25.0 ± 1.0 °C) was chosen to approximate ambient temperatures commonly experienced during the natural oviposition seasons of this species in the Osaka region. Approximately 40–60 individuals were reared in a plastic case (29.5 cm width, 19.0 cm depth, and 17.0 cm height) under long-day conditions (16-h light:8-h dark, 25.0 ± 1.0 °C) with a piece of moist cotton as a water source and oviposition site. They were fed an artificial diet (Masu Chigyo Super EPC-2, Marubeni Nisshin Feed Co., Ltd., Tokyo, Japan) and fresh carrot. The collected eggs were placed on a piece of moist cotton in plastic dishes (50 mm diameter, 12 mm depth) and continuously maintained under long-day conditions ^66^. For the experiments, first instar nymphs were kept in long-day conditions or were transferred to short-day conditions (12-h light:12-h dark, 25.0 ± 1.0 °C). Adult females reared under long-day conditions lay eggs destined to direct development (non-diapause), whereas those under short-day conditions lay eggs destined to enter diapause ^67^. In the present study, females reared under long- and short-day conditions are referred to as long- and short-day females, respectively. Eggs laid by long- and short-day females are referred to as long- and short-day eggs, respectively. Eggs deposited within a 2 h period were collected and used for experiments. These eggs were continuously maintained under long-day conditions until use for experiments. Under these conditions, the incidence of diapause was 14.6% in eggs laid by long-day females and 95.7% in eggs laid by short-day females, confirming that diapause fate was determined by maternal photoperiodic history rather than by environmental conditions experienced by the eggs.

### Genome and RNA sequencing

Alive *D. nigrofasciatus* was used to extract its genomic DNA. Total genomic DNA was extracted from 10 whole male bodies after removal of the digestive tract, as well as from the head and hind legs of a single male and female *D. nigrofasciatus*, using NucleoBond® HMW DNA (Macherey-Nagel, Germany) according to the manufacturer’s instructions. The resulting genomic DNA was size-selected using a Short Read Eliminator Kit (PacBio, CA, USA). Oxford Nanopore Technologies (ONT) sequencing libraries were then constructed and sequenced on the PromethION platform (Oxford Nanopore Technologies, UK) with the ligation sequencing kit SQK-LSK112 and flow cell R10.4. Base-calling was performed using Guppy basecalling software v6.3.8 with the model dna_r10.4_e8.1_sup. Finally, we obtained 30.40 Gbp ONT sequencing data; average and N50 read lengths were 19.11 Kbp and 31.18 Kbp, respectively. Additionally, using the same DNA sample, we prepared a whole genome sequencing library using the TruSeq DNA PCR-Free Library Prep Kit (Illumina, CA, USA) following the manufacturer’s protocol. This library was sequenced on the Illumina NovaSeq 6000 platform with 150 bp paired-end reads (PE150), generating 152.7 Gbp of short-read data. For chromosome-scale scaffolding by the Dovetail™ Omni-C™ Kit (Dovetail Genomics, CA, USA), the head and hind legs of another single male *D. nigrofasciatus* were used according to the manufacturer’s instructions. The Hi-C sequencing library was built on the Illumina NovaSeq 6000 platform with PE150 and generated 127.1 Gbp raw data. DNA purity and concentrations were measured by spectrometry using NanoPhotometer NP80-TOUCH (Implen, Germany) and fluorometry using Qubit 4 (Thermo Fisher Scientific, MA, USA).

Total RNA was extracted from whole eggs 12, 24, 40, 56, and 72 hours post-oviposition (hpo) using TRIzol Reagent with the PureLink RNA Mini Kit and PureLink DNase Set (Thermo Fisher Scientific) after homogenizing with the BioMasher II tissue grinder (Nippi, Inc., Tokyo, Japan) (Supplementary Table 16). Total RNA of 12, 24, 40, and 56 hpo eggs was purified with ssDNA / RNA Clean & Concentrator (ZYMO Research, CA, USA). RNA integrity was confirmed using a NanoDrop spectrophotometer (Thermo Fisher Scientific). For 12, 24, 40, and 56 hpo eggs, libraries were constructed using the NEBNext® Single Cell/Low Input RNA Library Prep Kit (New England Biolabs, MA, USA) and sequenced on an Illumina NextSeq 1000 platform with PE150. For 72 hpo eggs, libraries were prepared using the SMART-Seq® HT PLUS Kit (Takara Bio, Japan) and sequenced on an Illumina NovaSeq 6000 platform with PE150. To improve gene annotation and provide tissue-specific expression profiles, we also performed RNA-seq on adult tissues. Ovaries and neural tissues (brains and suboesophageal ganglions) were dissected from 10-day-old adult females at ZT 9-10, maintained under either long- or short-day photoperiod conditions. For these adult tissue samples, mRNA was enriched using the NEBNext® Poly(A) mRNA Magnetic Isolation Module, and libraries were prepared using the NEBNext® Ultra™ II Directional RNA Library Prep Kit (New England Biolabs). Both brain and ovary libraries were sequenced on an Illumina NovaSeq 6000 platform with PE150.

### Genome size estimation

Genome size was computationally estimated using k-mer frequency distribution analysis. Raw Illumina sequencing reads were processed using KMC v3 (K-mer Counter) ^68^ with k-mer length of 21. K-mers were counted. The resulting k-mer frequency histogram was generated using kmc_tools with a maximum coverage cutoff of 10,000. The k-mer frequency distribution was analyzed using GenomeScope2 ^69^ with k=21 and ploidy set to 2 (diploid).

Genome size was also estimated using flow cytometry to provide an independent validation of the computational estimates. The 1C (haploid) genome size of males and females of *D. nigrofasciatus* was estimated with reference to the method previously described ^70^. Uniform suspensions of single nuclei were extracted from a single *D. nigrofasciatus* head, a single *Drosophila virilis* female head (1C =328 Mbp) and a portion of brain from a *Periplaneta americana* male (1C =3300 Mbp) using the BD Cycletest Plus DNA Reagent Kit (BD Biosciences, CA, USA). After trypsin and RNase A treatments, the samples were incubated with propidium iodide for 10 min in the dark at 4 °C. Finally, the sample was filtered through a 50-μm nylon mesh. Cellular DNA content was assessed on the BD Accuri C6 Flow Cytometer (BD Biosciences). The obtained data were analyzed using the FCS Express software (De Novo Software, CA, USA). Haploid (1C) DNA quantity was calculated as (2C sample mean fluorescence/2C standard mean fluorescence) times 328 Mbp for the *D. virilis* standard and times 3300 Mbp for the *P. Americana* standard. The estimates based on the two standards were averaged to produce a 1C estimate for each sample.

### *De novo* genome assembly

Initial genome assembly was performed using MaSuRCA v4.1.0 ^71^ with a hybrid approach combining Illumina paired-end reads and Oxford Nanopore long reads. Illumina paired-end reads (153 bp mean insert size, 102 bp standard deviation) were provided as input along with the parameters FLYE_ASSEMBLY=0 to use CABOG assembly algorithm. The resulting primary assembly scaffolds were used for subsequent processing.

To remove redundant haplotigs from the assembly, we employed Purge Haplotigs v1.1.2 ^72^. Illumina paired-end reads were mapped to the primary assembly using minimap2 v2.24-r1122 ^73^ with the -ax sr option. Read-depth histogram was generated using the purge_haplotigs hist command to identify haploid and diploid peaks. Based on the histogram analysis showing an average coverage of 94.5×, cutoff values were set as: low=5, midpoint=70, high=190, with junk coverage threshold at 101. Coverage statistics were calculated using purge_haplotigs cov, followed by the actual purging step using purge_haplotigs purge.

Potential contamination in the assembly was removed using BlobTools v1.1.127 ^74^, which analyzes unexpected coverage, GC content, or similarity to bacterial and other contaminant sequences. Sequence coverage was determined by mapping Illumina reads with bwa v0.7.17-r1188. Similarity analysis was performed using BLASTn v2.13.0+ ^75^ against NCBI NT database v5 (options: -task megablast -culling_limit 10 -evalue 1e-25 -outfmt ‘6 qseqid staxids bitscore std sscinames sskingdoms stitle’). Mitochondrial genomes were identified through gene prediction using the MITOS2 webserver (accessed on November 28, 2023) and subsequently removed. As a result, one contig was removed as a bacterial genome and another as a mitochondrial genome.

The purged and decontaminated assembly was further scaffolded using Hi-C data. The input *de novo* assembly and Dovetail OmniC library reads were processed using HiRise ^76^, a software pipeline designed specifically for using proximity ligation data to scaffold genome assemblies.

Dovetail OmniC library sequences were aligned to the draft input assembly using bwa ^77^. The separations of Dovetail OmniC read pairs mapped within draft scaffolds were analyzed by HiRise to produce a likelihood model for genomic distance between read pairs. This model was used to identify and break putative misjoins, score prospective joins, and make joins above a threshold.

A Circos plot illustrating the distribution of genomic elements was generated using Circos v0.69-9 ^78^. The plot displays the major scaffolds arranged in a circular layout with tracks showing gene density, GC content, repeat element distribution, and other genomic features. The completeness of the genome assembly was evaluated using BUSCO (Benchmarking Universal Single-Copy Orthologs) v5.1.2 ^78^ implemented in gVolante ^79^. Assembly completeness was assessed against both the arthropoda_odb10 and insecta_odb10 lineage-specific ortholog datasets to evaluate the representation of conserved genes at different taxonomic levels.

### Prediction of repeat regions and annotation of protein-coding genes

For *de novo* repeat prediction, RepeatModeler v2.0.1 ^80^ was first used to identify novel repetitive elements in the genome. This *de novo* library was supplemented with known repeat sequences from closely related species to create a comprehensive repeat database. The combined repeat library was then used to identify and softmask repetitive elements in the genome using RepeatMasker v4.1.2 ^81^.

Structural annotation of protein-coding genes was performed on the soft-masked genome using three complementary approaches: *ab initio* prediction, homology-based prediction, and RNA-seq-based prediction. Each prediction method incorporated RNA-seq data from eggs, ovaries, and brains as supporting evidence.

The *ab initio* prediction was carried out using BRAKER v3.0.2 ^26–29,82,83^, incorporating protein data from OrthoDB 10’s arthropoda dataset and the mapping data of the filtered RNA-seq reads. This BRAKER prediction served as the foundation for our gene set. To complement and improve this base set, we employed two additional approaches. GeMoMa v1.9 ^29^ was used for homology-based prediction with gene sets from four model species (*Apis mellifera, Drosophila melanogaster*, and *Tribolium castaneum*), while StringTie2 v2.1.7 ^83^ was used for RNA-seq-based transcript assembly and gene prediction. The predictions from GeMoMa and StringTie2 were used to identify and add genes that BRAKER3 had missed. These combined predictions were merged and duplicate genes were removed using GffCompare v0.12.6 ^84^ with transcript classification codes set to u, i, and y to form a final, comprehensive consensus gene set.

Gene functional annotation was conducted using eggNOG-mapper online (http://eggnog-mapper.embl.de/) and BLASTp-based methods. For the BLASTp-based annotation, we searched against databases including *Homo sapiens* (GRCh38), *Mus musculus* (GRCm39), *Caenorhabditis elegans* (WBcel235), *Drosophila melanogaster* (BDGP6.32), and UniProt Swiss-Prot to identify the best hits for annotation (E-value < 1.0 × 10^−^5). The completeness of the genome assembly was also evaluated using BUSCO v5.1.2 ^78^ implemented in gVolante ^79^.

### Phylogenetic analysis

To reconstruct the phylogenetic relationships among cricket species, we analyzed protein sequences from eight species: *Acheta domesticus* ^85^, *Apteronemobius asahinai* ^86^, *Dianemobius nigrofasciatus, Gryllus bimaculatus* ^87^, *Laupala kohalensis* ^87^, *Locusta migratoria* ^88^, *Schistocerca gregaria* (GCF_023897955.1), and *Teleogryllus occipitalis* ^89^. Translated coding sequences (CDS) from each species were used as input for ortholog identification.

Orthologous gene groups were identified using OrthoFinder v2.5.4 ^90^ with default parameters. Single-copy orthologous genes (SCOs) were extracted for phylogenetic reconstruction, resulting in 334 SCOs shared across all eight species. Each SCO was aligned using MAFFT v7.505 ^91^ with the --auto option, which selected the L-INS-i strategy. The aligned sequences were subsequently trimmed using trimAl v1.4.rev15 ^92^ with the -automated1 option to remove poorly aligned regions and gaps.

The trimmed alignments were concatenated, and a partition file was generated to define gene boundaries within the concatenated alignment. Maximum likelihood phylogenetic analysis was performed using IQ-TREE v2.2.0.3 ^93^ with the following parameters: -bb 1000 for ultrafast bootstrap analysis with 1000 replicates, -alrt 1000 for SH-like approximate likelihood ratio test with 1000 replicates, -m MFP for automatic model selection using ModelFinder ^94^, and partition model optimization (-sp).

Divergence times were obtained from TimeTree (https://timetree.org/) ^95^ and mapped onto the resulting phylogeny to provide temporal context for the evolutionary relationships among the analyzed species.

### RNA-seq analysis

To prepare the reference transcriptome for quantification, the longest isoform for each gene was extracted from the gene models. Transcript abundance estimation was performed using Salmon v1.10.1 ^96^. First, a quasi-mapping index was built from the longest isoform cDNA sequences derived from the *D. nigrofasciatus* genome annotation. RNA-seq data from eggs collected at five developmental time points (12, 24, 40, 56, and 72 hpo) under both diapause (short-day) and non-diapause (long-day) conditions were analyzed. Paired-end reads were mapped using Salmon quant with automatic library type detection (-l A). Transcript-level estimates were imported and processed using the tximport package ^97^ in R. Transcripts per million (TPM) values and estimated read counts were extracted for downstream analysis.

Differential expression analysis was performed using DESeq2 v1.38.0 ^98^. Low-expressed genes were filtered by removing genes with total read counts less than 10 across all samples. Library size normalization and dispersion estimation were performed using default DESeq2 parameters, followed by the Wald test for differential expression. Results were ordered by adjusted *p*-value (Benjamini-Hochberg correction) and exported for further analysis.

Gene Ontology (GO) and KEGG pathway enrichment analyses were performed using g:Profiler (https://biit.cs.ut.ee/gprofiler/) ^99^. Differentially expressed genes were mapped to their *Drosophila melanogaster* orthologs based on the BLASTp annotation results, and the *D. melanogaster* gene IDs from the BDGP6.32 reference genome were used as input for enrichment analysis.

### Principal component analysis

To visualize the overall transcriptional differences between diapause and non-diapause conditions across developmental time points, three-dimensional principal component analysis (PCA) was performed. TPM values from all samples were used. Genes with zero variance across all samples were removed prior to analysis. PCA was performed using the prcomp function in R with scaling (scale. = TRUE).

The first three principal components were extracted and visualized using the rgl package for 3D plotting. Samples were colored according to their diapause status (diapause vs. non-diapause) to assess the separation of conditions in the reduced dimensional space. The interactive 3D visualization was generated using plot3d functions.

### Time-series clustering analysis, cluster comparison and classification

To identify genes with similar temporal expression patterns during embryonic development, time-series clustering was performed using the Mfuzz package ^98^. First, normalized count data from DESeq2 were obtained for each developmental time point (12h, 24h, 40h, 56h, and 72h). The mean expression values were calculated separately for diapause and non-diapause conditions, and genes with zero standard deviation were removed.

To focus on genes with substantial expression changes across time points, genes with a coefficient of variation less than 1.0 were filtered out. This filtering step ensures that the clustering analysis captures biologically meaningful expression dynamics rather than noise from low-variance genes.

The filtered expression data were converted to ExpressionSet objects and standardized using the Mfuzz standardize function. The fuzziness parameter (m) was estimated using the mestimate function. Based on the elbow method analysis and practical considerations, the number of clusters was set to 20 for both diapause and non-diapause datasets.

To identify relationships between diapause and non-diapause expression patterns, cosine similarity was calculated between all pairwise combinations of cluster centroids. Cluster pairs were classified as “homochronic” (similar temporal dynamics) or “heterochronic” (distinct temporal dynamics) based on a cosine similarity threshold of 0.8. This classification allows for the identification of genes that maintain similar expression patterns across conditions versus those that show condition-specific temporal regulation. Gene overlap significance between clusters was assessed using upper-tail hypergeometric tests. Cluster pairs with *p*-values < 10^-10^ were considered significantly overlapping, indicating non-random association of genes between diapause and non-diapause conditions.

For functional interpretation, genes from cluster pairs in the top and bottom 10th percentiles of cosine similarity were extracted. These gene sets, representing the most homochronic and heterochronic patterns respectively, were subjected to GO and KEGG enrichment analysis using g:Profiler ^99^ to identify biological processes associated with conserved versus divergent temporal expression dynamics during diapause induction.

### ATAC-seq library preparation and sequencing

Chromatin accessibility profiling was performed using ATAC-seq on 24 hpo eggs. For either long- or short-day conditions, two biological replicates were prepared (Supplementary Table 17).

Libraries were prepared using the ATAC-seq Kit (Active Motif, CA, USA). Frozen egg samples were lysed in 100 μL ice-cold ATAC Lysis Buffer and mechanically disrupted using scissors to release nuclei. Cell debris and eggshell fragments were removed by filtration through pluriStrainer-Mini 40 μm filters (pluriSelect, Germany). Subsequent library preparation steps were performed according to the manufacturer’s protocol.

Library quality was assessed using NanoDrop One (Thermo Fisher Scientific) for concentration measurement and DNA ScreenTape (Agilent Technologies, CA, USA) for fragment size distribution analysis. Qualified libraries were sequenced on an Illumina NovaSeq 6000 platform with 150PE. The sequencing generated 11.15-11.60 Gbp of data per sample.

### ATAC-seq data analysis

Raw sequencing reads were quality-trimmed using Skewer v0.2.2 ^100^ with the following parameters: maximum allowed error rate of 0.1, maximum allowed indel error rate of 0.03, and minimum read length of 18 bp after trimming. Paired-end mode was enabled for all samples.

Trimmed reads were aligned to the *D. nigrofasciatus* reference genome using Bowtie2 v2.3.4.2 ^101^ with parameters -N 1 (maximum mismatch of 1) and -X 2000 (maximum fragment length of 2000 bp). Peak calling was performed using MACS2 v2.1.2 ^102^ with a q-value threshold of 0.05 (- q 0.05) and parameters --nomodel --shift -75 --extsize 150, which defines a 150 bp region centered on the read start position.

To create a unified peak set, detected peaks from all samples were sorted and merged using bedtools v2.27.1 ^103^ with default settings. The resulting consensus peak regions were formatted into SAF format for downstream analysis.

Genomic annotation of peaks was performed by categorizing each peak into the following genomic regions: 1-5 kb upstream (1to5kb), <1 kb upstream (promoter), 5’UTR, exon, intron, 3’UTR, exon/intron boundary (defined as 100 bp surrounding exon-intron junctions), and intergenic regions. When peaks overlapped multiple features, priority was given in the following order: 5’-/3’-UTR > exon/intron boundary > exon/intron. Genes located within 1 Mbp of each peak were assigned as nearby genes.

Read counts for consensus peaks were quantified using featureCounts v1.6.3 ^104^. Differential accessibility analysis between diapause-inducing and non-diapause conditions was performed using edgeR v3.26.8 ^105^ with TMM normalization. Log2 fold changes were calculated for each peak region. Peaks with uncorrected *p*-values < 0.05 were considered differentially accessible, and their associated nearby genes from different genomic contexts were extracted for functional analysis. Genomic contexts included: 1-5 kb upstream regions (1-5 kbp upstream of transcription start sites, TSS), promoter regions (<1 kbp upstream of TSS), 5’UTR, exon, intron, 3’UTR, exon/intron boundaries (within 100 bp of exon-intron junctions), and intergenic regions (areas not classified into any of the above categories).

## Data availability

Raw read data used in this article (RNA-seq, ATAC-seq, Omni-C and WGS) have been submitted at DNA Data Bank of Japan (DDBJ) repository under accession no. PRJDB18895 and are publicly available as of the date of publication. The datasets are available in following link; https://ddbj.nig.ac.jp/resource/bioproject/PRJDB18895. The assembled genome and annotation datasets are available in https://doi.org/10.6084/m9.figshare.29665742.v1. The scripts used for the analyses in this study are available on GitHub (https://github.com/Kataoka-K-Lab/Dnigrofasciatus_EggDiapause.git).

## Acknowledgments

This work was supported by the Cooperative Research Grant of the Genome Research for BioResource (NODAI Genome Research Center, Tokyo University of Agriculture), Cross-ministerial Moonshot Agriculture, Forestry and Fisheries Research and Development Program, “Technologies for Smart Bio-industry and Agriculture” (Bio-oriented Technology Research Advancement Institution, BRAIN) [Grant Number: JPJ009237], JSPS KAKENHI Grant-in-Aid for Scientific Research (C) [Grant Number: 21K05614], and Grant-in-Aid for JSPS Fellows [Grant Number: 21J23478/22KJ2609]. We thank Dr. Tarô Adati (Department of International Agricultural Development, Faculty of International Agriculture and Food Studies, Tokyo University of Agriculture) for his valuable advice and support.

## Author Contributions

K.K., Y. S., and S.G.G. designed research; K.K., Y.S., R.S., Y.K., K.N., K.Y., and T.A. performed research; K.K. analyzed data; and K.K., Y.S., and S.G.G. wrote the paper.

## Competing Interest Statement

The authors have no competing interests to disclose.

## Notes

### Competing Interest Statement

The authors have declared no competing interest.

### Summary of Updates

Discussion revised; Figure1, 2, 4 revised

