## Supplementary material for "Temporal orchestration of transcriptional and epigenomic programming underlying maternal embryonic diapause in a cricket model": Fig. S1-12

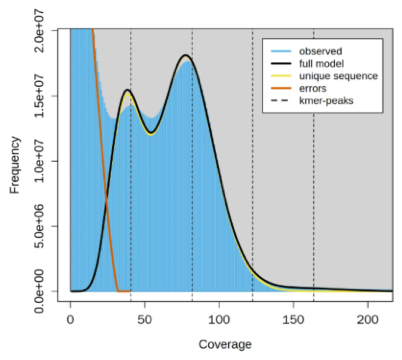

Fig. S1. *k*-mer frequency analysis using GenomeScope2 (*k*=21) with Illumina short reads. GenomeScope2 profile showing a heterozygosity peak based on k-mer distribution analysis. The x-axis represents coverage, and the y-axis shows the frequency.

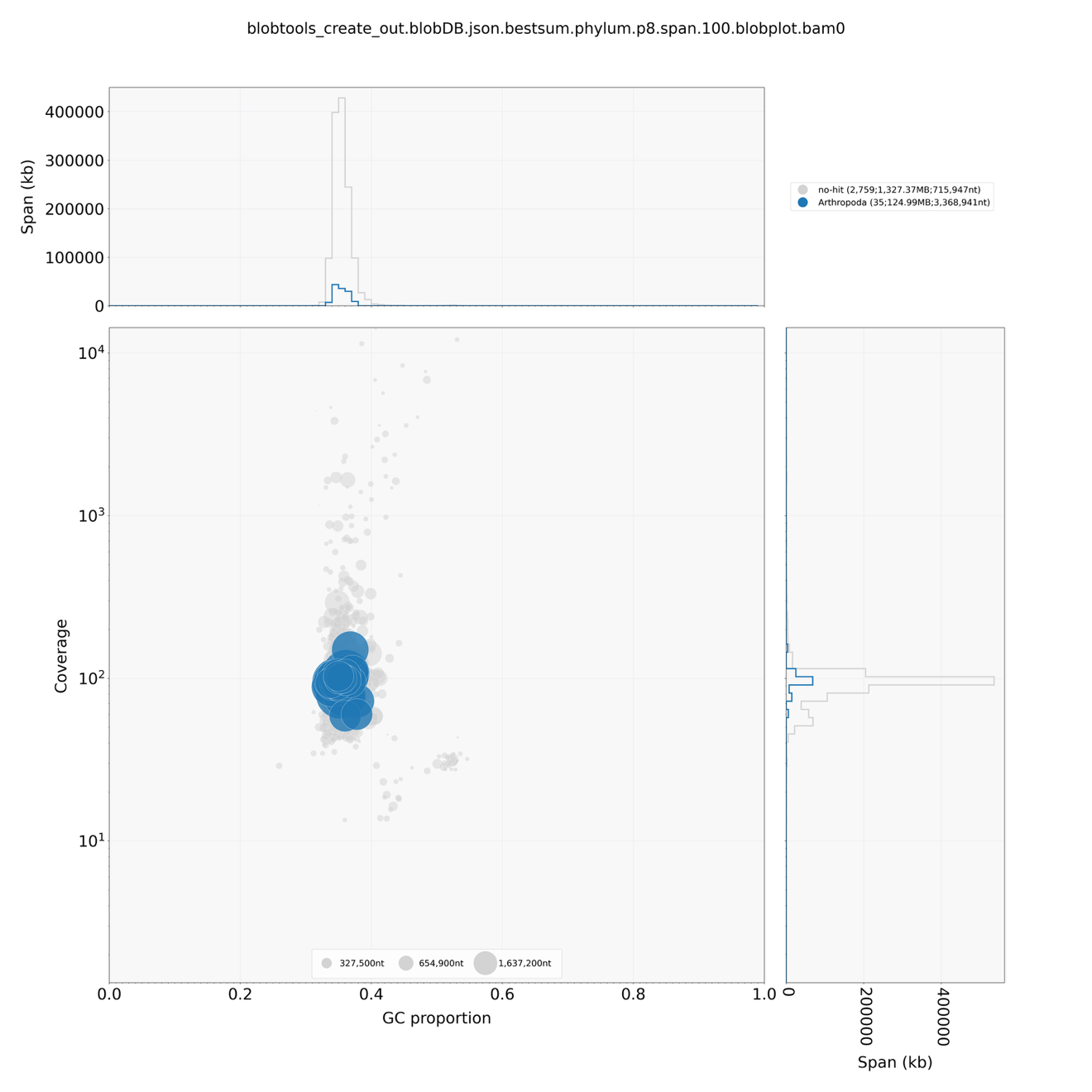

Fig. S2. BlobTools analysis of the *D. nigrofasciatus* contig-level *de novo* genome assembly. Assessment of assembly quality and potential contamination using BlobTools. The main scatter plot shows GC content proportion (x-axis) versus sequence coverage (y-axis, log scale) for each contig in the assembly. Point size represents contig length (327.5 kb, 654.9 kb, and 1.64 Mb shown in legend). Blue points indicate contigs identified as Arthropoda, while gray points represent sequences with no taxonomic assignment. The top histogram shows the distribution of contigs by span (kb), and the right histogram shows the distribution by coverage.

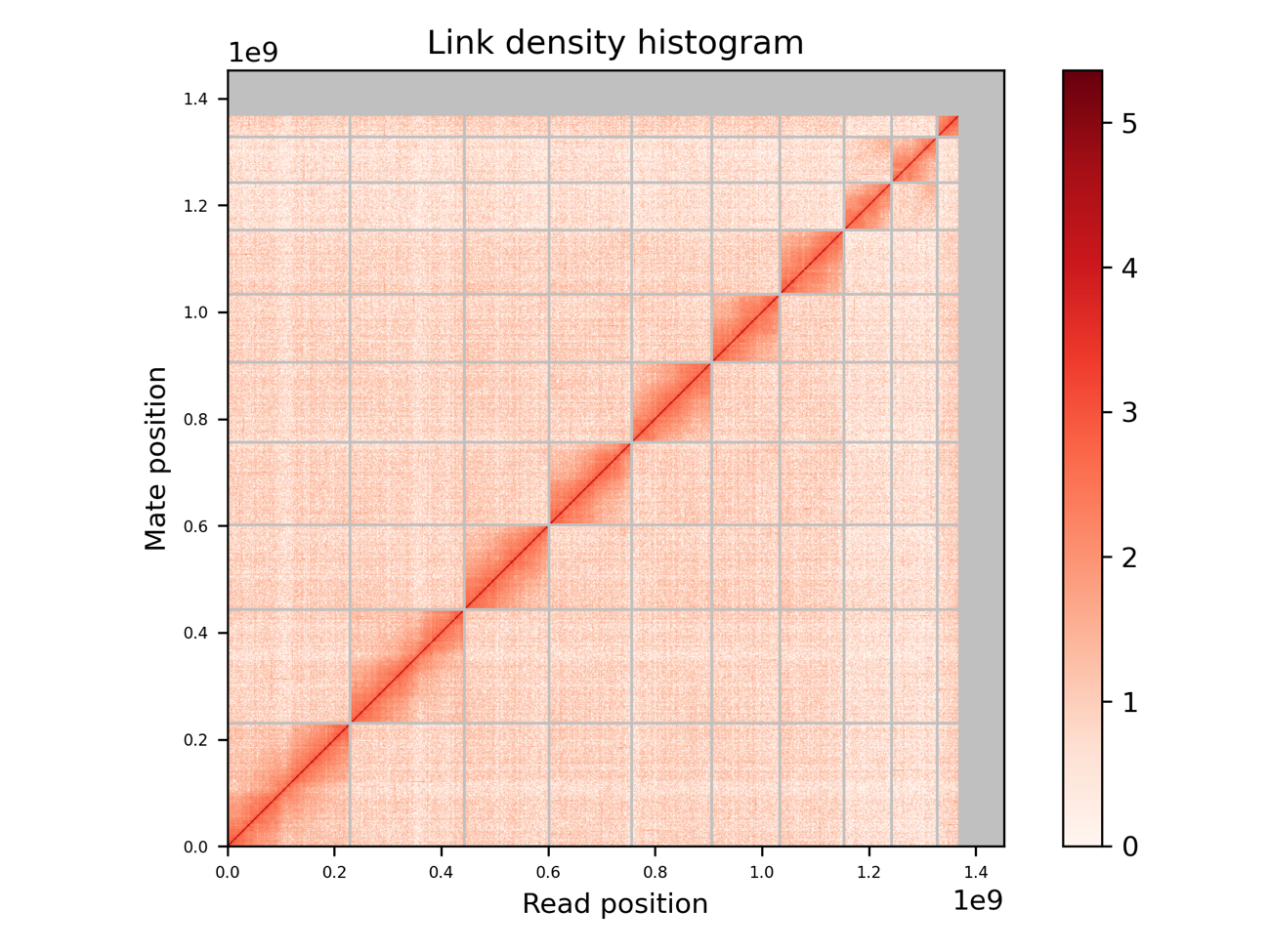

Fig. S3. Hi-C chromosome contact map.

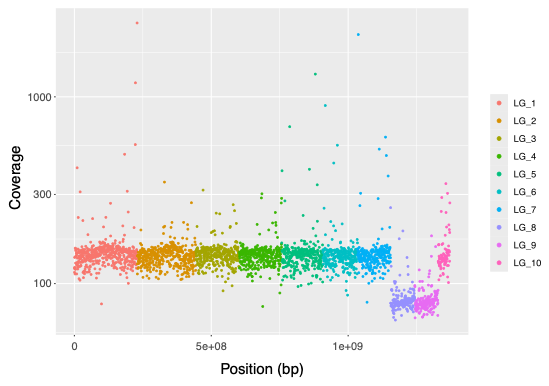

Fig. S4. Identification of X-linked scaffolds through male genomic read coverage analysis. Coverage depth analysis of male genomic DNA reads mapped across the ten linkage groups (LG1-10) of *D. nigrofasciatus*. Each point represents the mean coverage in a 500 kb window calculated using mosdepth. The x-axis shows the position along each linkage group, and the y-axis displays the coverage depth. LG8 (purple) and LG9 (pink) exhibit approximately half the coverage (~50×) compared to autosomal linkage groups (~100×), indicating their X-linked nature in the XO male sex determination system. The reduced coverage of these two linkage groups is consistent with males carrying a single copy of the X chromosome, while autosomes are present in two copies. Together, LG8 and LG9 likely represent the complete X chromosome, which failed to assemble into a single scaffold possibly due to repetitive sequences at their junction.

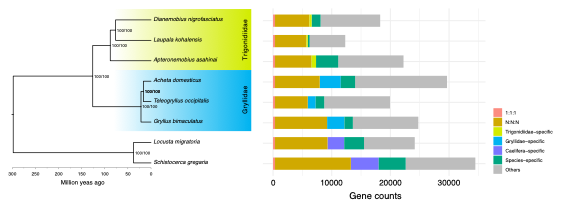

Fig. S5. Estimated phylogenetic tree and gene contents in Orthoptera. Maximum likelihood phylogenetic tree of orthologous single-copy genes from eight Orthopteran species (left) with node support values and gene family distribution (right). Node values represent SH-aLRT/Ultrafast bootstrap supports in percentage. The time scale is millions of years ago (Mya). Bar plots show the distribution of gene categories: 1:1:1 orthologs (pink), N:N:N orthologs (golden brown), Trigonidiidae-specific (light green), Gryllidae-specific (blue), Caelifera-specific (purple), species-specific (green), and others (gray).

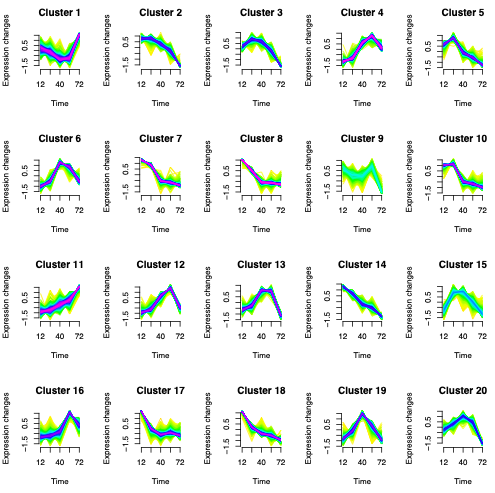

Fig. S6. Temporal expression patterns of gene clusters in long-day eggs identified by fuzzy c-means clustering. Fuzzy c-means clustering analysis of time-series transcriptome data from long-day eggs, showing 20 distinct temporal expression patterns across five developmental time points (12, 24, 40, 56, and 72 hours post-oviposition). Each panel represents a cluster with normalized expression changes (y-axis) plotted against time (x-axis). Individual gene expression profiles are shown as colored lines, with color intensity representing membership values in the fuzzy clustering approach. The blue line indicates the cluster centroid representing the average expression pattern.

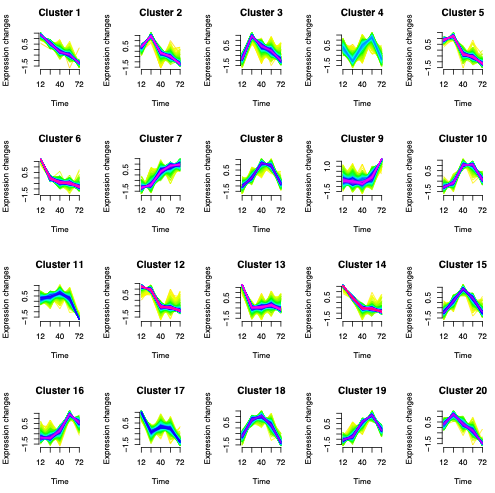

Fig. S7. Temporal expression patterns of gene clusters in short-day eggs identified by fuzzy c-means clustering. Analysis of time-series transcriptome data from short-day eggs, revealing 20 distinct temporal expression patterns across the same five developmental time points (12, 24, 40, 56, and 72 hours post-oviposition). Cluster organization and visualization are as described in Fig. S5. In contrast to non-diapause eggs, these expression patterns reflect the transcriptional programs associated with preparation for and entry into embryonic diapause at the cellular blastoderm stage.

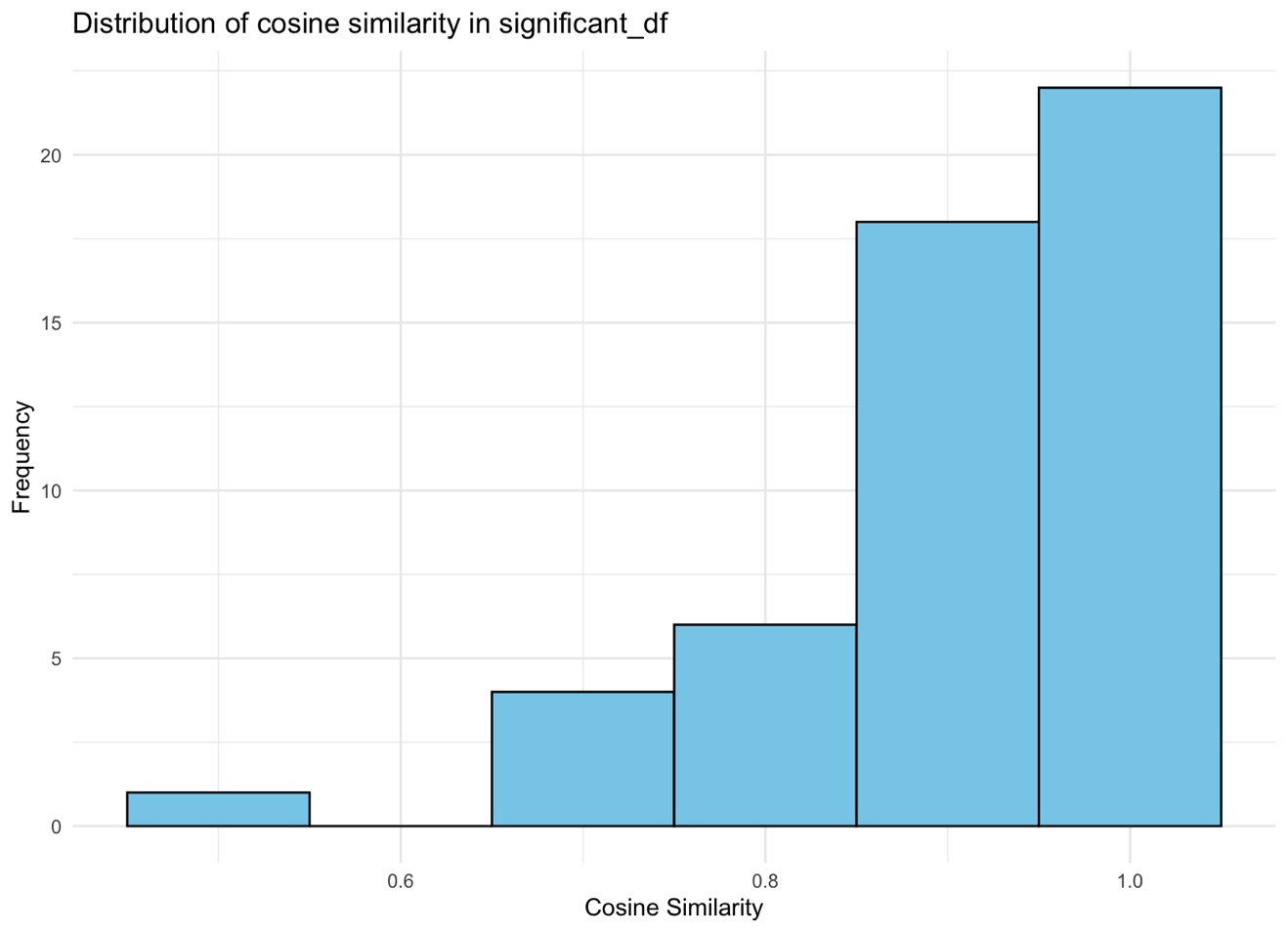

Fig. S8. Distribution of cosine similarity values between significantly overlapping cluster pairs (long- vs. short-day eggs). Histogram showing the frequency distribution of cosine similarity values calculated between cluster centroids from long- and short-day egg transcriptomes. Only cluster pairs with significant gene overlap (hypergeometric test, *p*-value < 10⁻¹⁰) are included in this analysis.

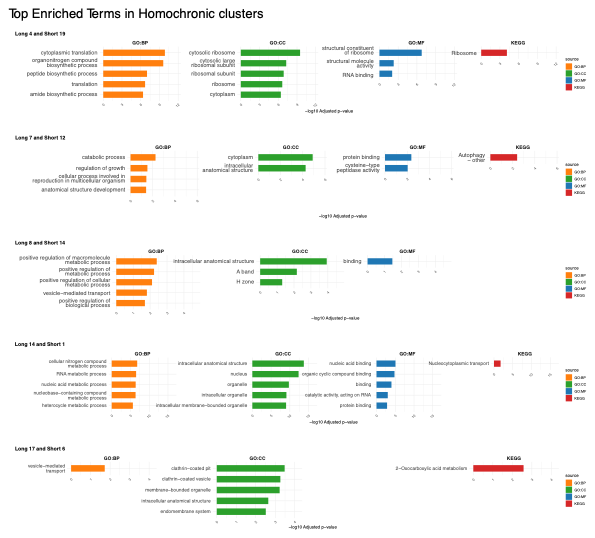

Fig. S9. Gene Ontology and KEGG pathway enrichment analysis of the top five homochronic cluster pairs. Functional enrichment analysis of genes from the five cluster pairs with the highest cosine similarity values between long- and short-day eggs. Each panel represents one homochronic cluster pair (Long_X and Short_Y format indicates non-diapause and diapause clusters, respectively). Bar plots show the top enriched terms for GO Biological Process (GO:BP, orange), GO Cellular Component (GO:CC, green), GO Molecular Function (GO:MF, blue), and KEGG pathways (red). The x-axis represents -log10 adjusted *p*-values, with longer bars indicating more significant enrichment.

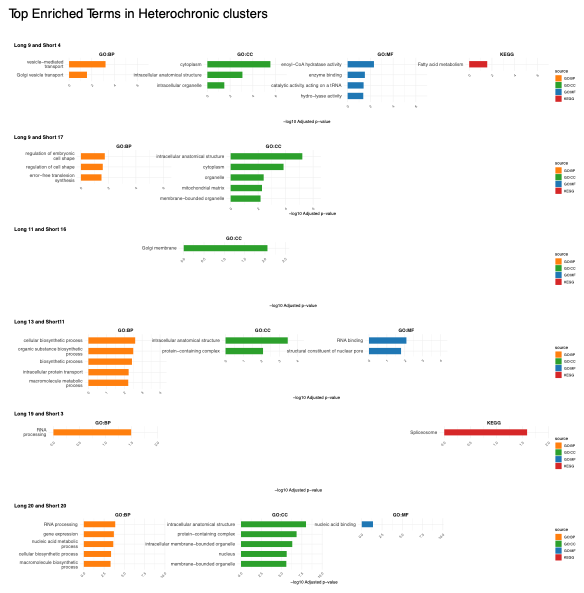

Fig. S10. Gene Ontology and KEGG pathway enrichment analysis of the top five heterochronic cluster pairs. Functional enrichment analysis of genes from the six cluster pairs with the lowest cosine similarity values between long- and short-day eggs. Panel organization and visualization are as described in Fig. S8, with each panel showing one heterochronic cluster pair. These clusters represent genes with divergent temporal expression patterns between diapause-destined and non-diapause eggs, highlighting biological processes that undergo condition-specific temporal regulation during early embryonic development.

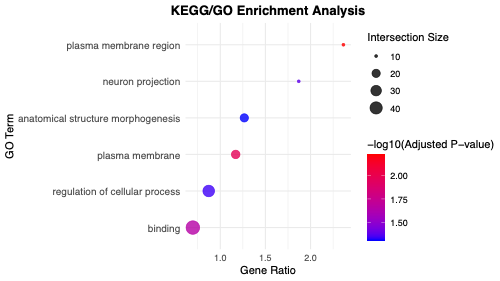

Fig. S11. Gene Ontology enrichment analysis of genes associated with decreased chromatin accessibility in short-day eggs. Functional enrichment analysis was performed using genes located near chromatin regions showing reduced accessibility in short-day eggs compared to long-day eggs, as identified by ATAC-seq. The x-axis shows the gene ratio (percentage of query genes in each GO term), and the y-axis displays the top enriched GO terms across biological process (BP), molecular function (MF), and cellular component (CC) categories. Dot size represents the number of genes in the intersection between the query and the GO term (intersection size), while color intensity indicates the statistical significance (-log10 adjusted *p*-value).

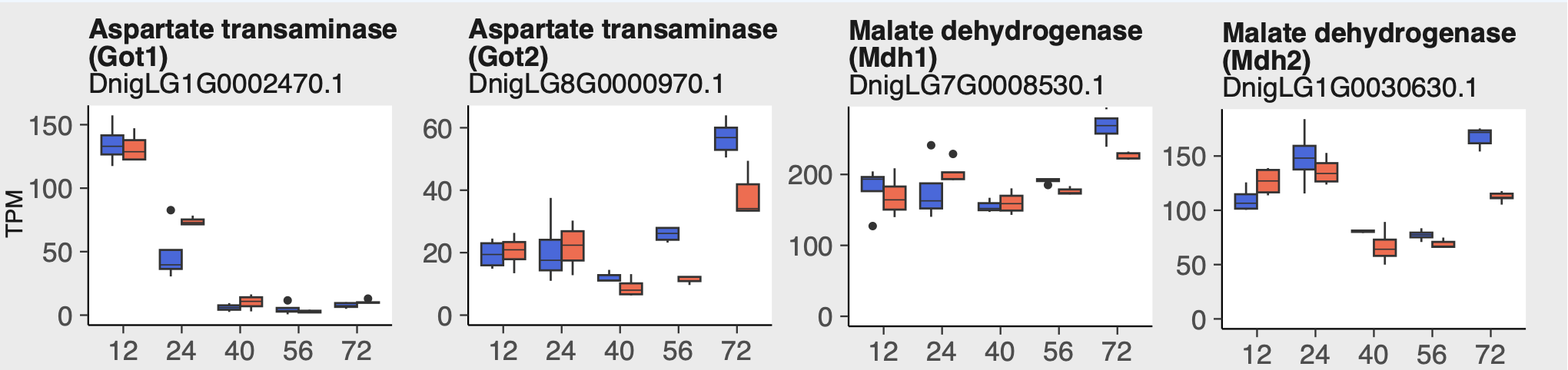

Fig. S12. Temporal expression profiles of gluconeogenic enzyme genes in long- and short-day eggs. Expression patterns of aspartate transaminase (*Got1* and *Got2*) and malate dehydrogenase (*Mdh1* and *Mdh2*) genes across developmental time points (12, 24, 40, 56, and 72 hours post-oviposition). Blue boxes represent short-day eggs and red boxes represent long-day eggs. Data are shown as box plots with TPM (Transcripts Per Million) values on the y-axis. These enzymes are involved in the malate-aspartate shuttle and gluconeogenesis pathways, facilitating the conversion of amino acid-derived carbon skeletons to glucose precursors.
